# Bacterial mobbing behavior: coordinated communal attack of *Pseudomonas aeruginosa* on a protozoan predator

**DOI:** 10.1101/2020.06.15.152132

**Authors:** N. Shteindel, Y. Gerchman

## Abstract

Mobbing, a group attack of prey on predator, is a strategy enacted by many animal species. Here we report bacterial mobbing carried out by the bacterium *Pseudomonas aeruginosa* towards *Acanthamoeba castellanii*, a common bacterivore. This behavior consists of bacterial taxis towards the amoebae, adhesion en masse to amoebae cells, and eventual killing of the amoebae. Mobbing behavior transpires in second’s timescale and responds to predator population density. A mutant defective in the production of a specific quorum sensing signal displays reduced adhesion to amoeba cells. This deficiency ameliorated by external addition of the missing signal molecule. The same quorum sensing mutant also expresses long term deficiency in its ability to cause amoeba death and shows higher susceptibility to predation, highlighting the importance of group coordination to mobbing and predation avoidance. These findings portray bacterial mobbing as a regulated and dynamic group behavior.

## Introduction

Mobbing is a predation avoidance behavior, manifested as an attack on predator by a group of prey organisms[1]. Predation by bacterivores is a major selective force shaping bacterial evolution, leading to the development of many predation avoidance mechanism - increasing of size, either per cell or by microcolony formation, anti-predator toxin secreation and surface signal masking[2, 3]. Nevertheless, protozoan predation can be a fast process, with several to several thousand bacteria consumed every minute[4, 5], making these slow mechanisms of limited effectivity. Mobbing behavior seem to be a natural direction for bacterial evolution, as they often live in large clonal populations and able to communicate via Quorum Sensing (QS)[6]. Still, no case of bacterial mobbing was reported to date. *Pseudomonas aeruginosa* is a common and ubiquitous bacterium known for its communal adaptations[7], that was shown to kill *Acanthamoeba castellanii*, a common soil bacterivore in co-culture[8]. Here we study the interaction of these two organisms in seconds and minute’s timescale, showing that the killing of amoebae is the product of a fast and direct communal attack behavior - mobbing.

## Methods

### Strains, plasmids and culture conditions

#### Pseudomonas aeruginosa

PAO1 w.t and *P. aeruginosa* PAO1 Δ*pqs*A[9] carrying the pMRP9-1 plasmid[10, 11], encoding for Carbenicillin resistance and constitutive expression of GFP_mut2_ (ref. 11) were cultivated in 50 ml M9 medium (47.75 mM Na_2_HPO_4_, 22.05 mM KH_2_PO_4_. 8.56 mM NaCl, 18.69 mM NH_4_Cl, 2 mM MgSO_4_, 0.1 mM CaCl_2_), 200 µg/ml Carbenicillin, 0.4% glucose in 100 ml Erlenmeyer flask, in 37° C, 120 RPM for a period of 18 hours, centrifuged to separate (12,000 g, 1 minute), washed once in and re-suspended TBSS in a TRIS-buffered salts solution (TBSS) (2 mM KCl, 1 mM CaCl2, 0.5 mM MgCl2, 1 mM TRIS).

#### Acanthamoeba castellanii

was cultured in PYG medium (ATCC 712) supplemented with 100 µg/ml Gentamicin, 10 ml in a 50 ml tissue culture treated culture flasks (Greiner, Germany) in 25°C, static, for five days. The flask was shaken vigorously to separate the cells from the plastic surface; culture was transferred to 1.5 plastic micro tubes (1.5 ml per tube) and centrifuged to separate the cells (200 g, 30 sec) The culture was gradually transferred to TBSS medium, replacing 500, 1000, 1500 μL of the medium in each tube in consecutive wash cycles, centrifuged once more and collected into 1 ml of TBSS in a 10 mm glass tube. Culture density was determined by microscopy in a disposable penta-square counting chamber (Vacutest Kima, Italy) and diluted to the culture density indicated in each experiment.

### Microscopy of *P. aeruginosa* PAO1 attachment to amoeba

Ten µL of 5X10^5^ cells/ml amoeba culture in TBSS medium were added to a counting chamber and imaged using a fluorescence enable binocular system (Nikon SMZ18 fluorescence dissecting microscope connected to a Nikon DS-Fi3 camera, using the NIS elements software) in visible light and in green fluorescence. Fluorescence imaging setting: magnificationX12, exposure time 500 milliseconds, gain X14, field size 2880X2048 pixels, dynamic range of 3X8 bit. In this setting, using a plasmid that produces mild fluorescence, only aggregated bacteria can be seen. After imaging the amoebae in the absence of bacteria for a few minute, 10 µL of fluorescently tagged *P. aeruginosa* PAO1 culture were added and photographed every 10 seconds for a period on 10 minutes.

### Effect of amoeba population density on *P. aeruginosa* attachment behavior

Bacterial adhesion to amoebae was quantified using a kinetic assay in 96 well plate format[12] (figure 2a of this work). Amoeba culture in TBSS was diluted to 8X10^4^ cells/ml and 50 µL were pipetted into the first row of a 96 well plate (clear tissue culture treated polystyrene, flat bottom, Jet-biofil, China), and diluted in a double dilution series using fresh TBSS. The 12^th^ column (no amoeba) was added with only 50 µL TBSS. Amoebae were left to settle on the plate bottom for one hour prior to the addition of bacteria. Overnight culture of *P. aeruginosa* PAO1 was washed three times with TBSS, OD_600nm_ adjusted to 0.1 (measured in 100 µL volume in a clear flat bottom 96 well plate) and supplemented with 1.6 mg/ml Red#40 dye (Sigma, Israel). Fifty µL of this culture were pipetted to rows A-G of the plate containing the amoeba, row H pipetted with 50 µL of TBSS supplemented with the dye to be used as blank. Pipetting of bacterial culture to the plate was carried out within 30 seconds, using an 8-channel pipetor. Final bacterial culture density was OD_600nm_=0.05, final dye concentration 0.8 mg/ml and final amoebae counts 0, 4, 8, 16, 32, 64, 125, 250, 500, 1 000, 2 000 and 4 000 amoebae per well. The plate was loaded into a multimode plate reader (Synergy HT, Biotek, USA) and read kinetically for bottom fluorescence (Excitation 485_nm_/20, Emission 528_nm_/20, Gain 60) for one hour in one minute intervals.

**Figure 1.**
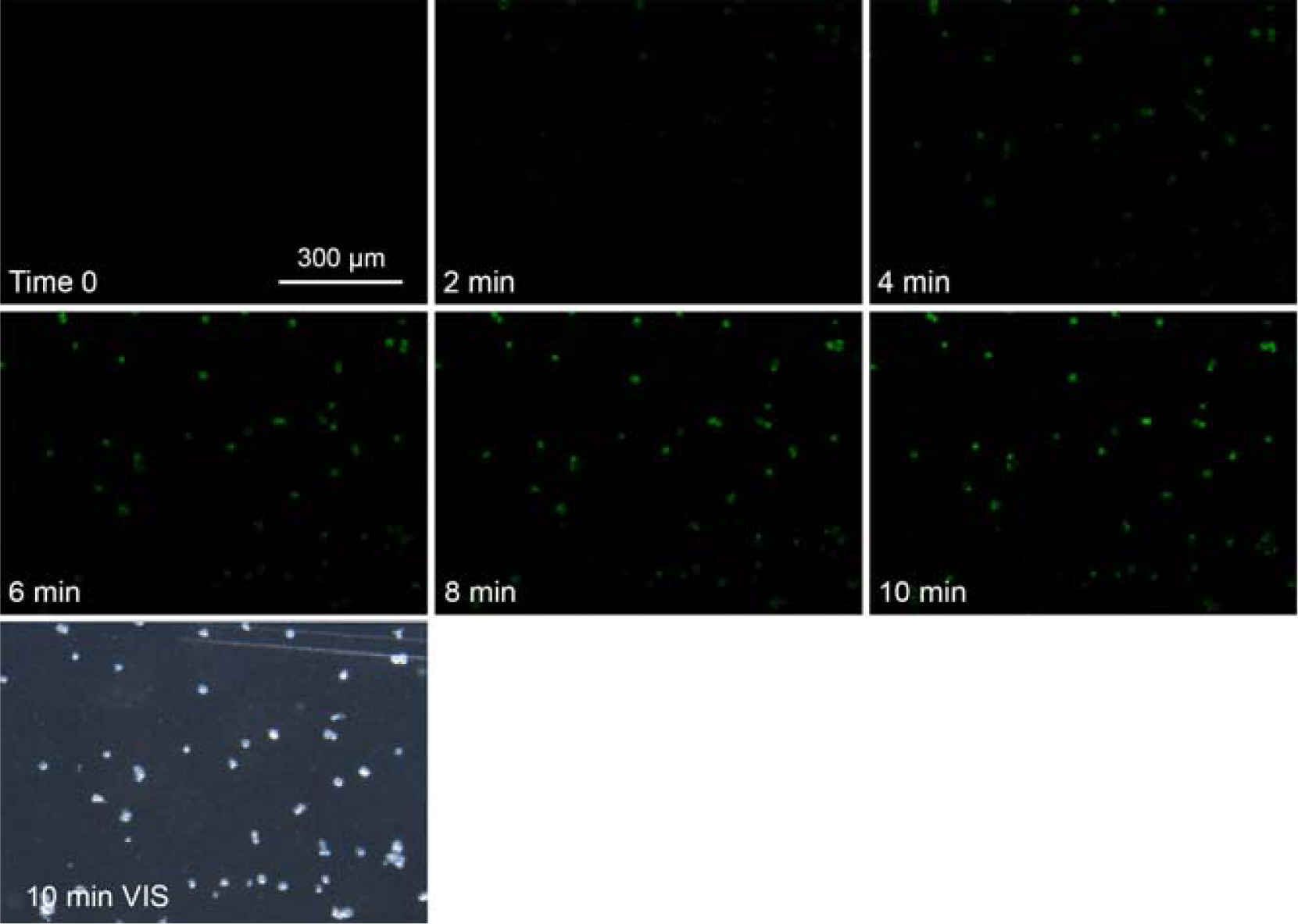
Time laps of GFP expressing *P. aeruginosa* adhesion to cells of *A. castellanii*. Fluorescence intensity rise as bacteria aggregate on amoebae cells. Plateau is reached within 10 minutes.

**Figure 2.**
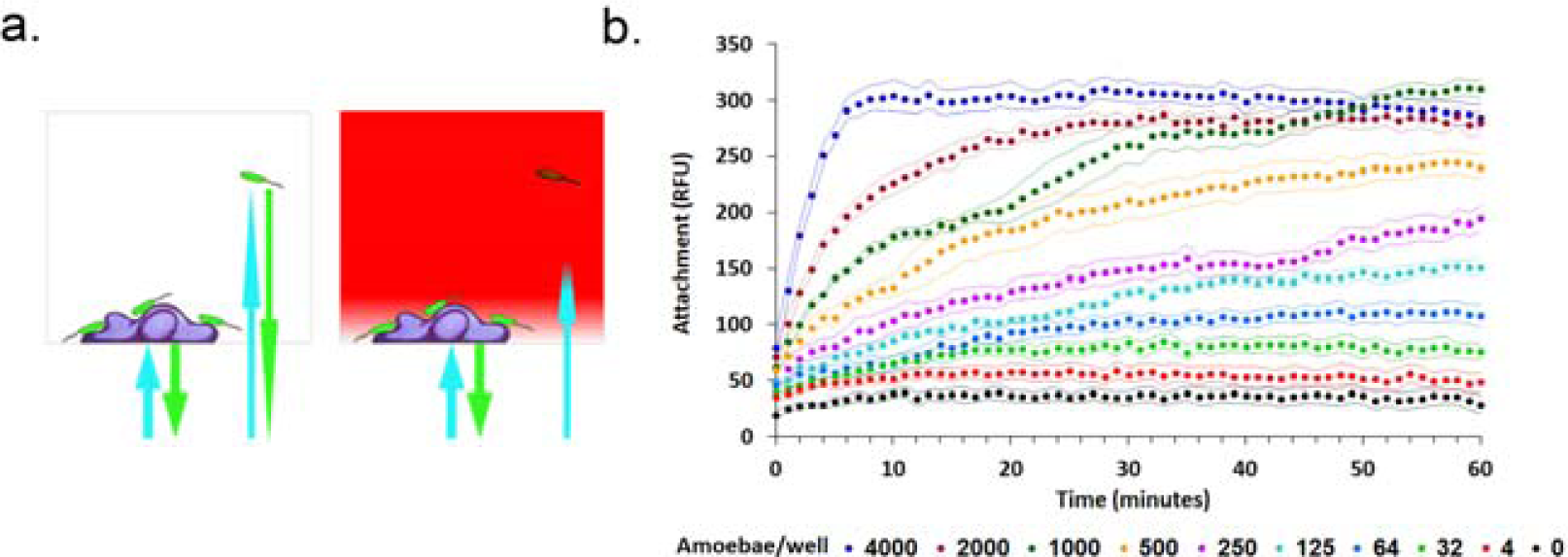
The effect of amoebae population density on *P. aeruginosa* adhesion kinetics. a. Illustration of bacterial adhesion kinetics in micro plate assay: fluorescent signal in the absence (left) and presence (right) of dye. Addition of the dye limits the depth of field to about 5 µm from the bottom of the well, allowing the detection of adhered GFP expressing bacteria to the well bottom in real time. b. Adhesion kinetics of *P. aeruginosa* to amoeba on the bottom of the microtiter wells. n=7 per for all treatments, dots signify measurements, flanking curves stand for ±1 S.D.

### *P. aeruginosa* taxis towards amoeba

Taxis experiments were done using Corning® FluoroBlok™ HTS 24-well Multi-well Permeable Support System with 3.0 µm high density PET intervening membrane (Corning, New York, USA) designed for cell migration assays. *Pseudomonas aeruginosa* and amoeba cultures were grown and prepared as described. Bacteria were diluted in TBSS to OD_600nm_=0.1 and amoebae culture was diluted to 1X10^4^ cells/ml. Amoeba culture (750 µL) were added to the bottom chamber of columns 1-3 of the plate, and columns 4-6 were added with 750 µL of TBSS buffer. The filter system was mounted onto the plate base and the plate was loaded onto the plate reader. The plate was read for bottom fluorescence (Excitation 485nm/20, Emission 528nm/20, gain 60, used as blank reading) to obtain the base fluorescence without bacteria. Then top chambers were loaded, one at a time, with 100 µL of the bacterial culture and read kinetically for bottom fluorescence every four seconds for a period of two minutes - appearance of fluorescence indicating the migration of bacteria from the upper chamber, through the membrane, to the bottom chamber.

### Modulation of *P. aeruginosa* adhesion behavior by amoebae conditioned buffer

Adhesion of *P. aeruginosa* w.t. in the presence and absence of A. castellanii conditioned buffer was carried using kinetic assay in 96 well plate format as previously described. Amoebae were culture and washed as described earlier, diluted to 10 000 cells per ml in TBSS medium and incubated in 25° C for 2 hours and separated by centrifugation (200 g, 1 min). Buffer separated from amoebae culture and unconditioned buffer were pipetted into 96 well plate in 50 µL volume. Fifty µL of w.t. PAO1 suspension, supplemented with RED#40 prepared was added to each well, and bottom fluorescence was read kinetically as described earlier.

### Effect of Pseudomonas quinolone signal (PQS) signaling on *P. aeruginosa* predation

Amoebae were cultured and transferred and diluted to 2X10^4^ amoebae/ml as previously described. Fifty µL of this culture (1,000 amoeba per well) were pipetted to 48 wells of a flat bottom clear 96 well plate, the other half pipetted with sterile TBSS. GFPmut2 expressing *P. aeruginosa* PAO1, either w.t. or Δ*pqs*A, were cultured, washed and diluted to OD_600nm_=0.1 as described earlier, and added into wells with or without amoeba (14 replicates per treatment). Bottom fluorescence reading (Excitation 485_nm_/20, Emission 528_nm_/20, Gain 60) was taken every 30 minutes over a period of 27 hours in order to assess bacterial population density kinetics.

### Effect of PQS concentration on *P. aeruginosa* attachment to amoebae

*Pseudomonas aeruginosa* PQS (2-nonyl-3-hydroxy-4-Quinolone, Sigma, Israel) was dissolved in DMSO to 10 mM concentration, diluted in TBSS medium in double dilution series, to concentrations of 20µM to 20 nM per 96-well plate well (11 concentrations + negative control; 25 µL volume). Amoebae culture was prepared as previously described, diluted to 4,000 cells/ml, and added to all above wells, 25 µL and 1,000 cells per well. GFP_mut2_ expressing *P. aeruginosa* PAO1 Δ*pqs*A was cultivated and prepared as described earlier, diluted to OD_600nm_=0.1 in TBSS and supplemented with 1.6 mg/ml Red#40. Fifty µL of this bacterial culture was added to rows A-G of the amoeba-PQS plate, to final volume of 100 µL, culture density of OD_600nm_=0.05, dye concentration of 0.8 mg/ml, 1,000 amoebae per well and 5 µM to 5 nM of PQS. Row H was added with dye supplemented TBSS and used as blanks. The plate was loaded to the plate reader and read for bottom fluorescence (Excitation 485_nm_/20, Emission 528_nm_/20, Gain 60) every minute for a period of one hour. The same experiment was conducted in the absence of amoebae (replaced with additional 25 µL of TBSS per well).

### Effect of PQS signaling on amoebae killing

PAO1 w.t. and Δ*pqs*A were cultured overnight in M9 medium. *E. coli* DH5α was cultured in Lennox LB (Himedia, Mumbai, India). All strains were washed twice in M9 buffer, and diluted to OD_600nm_= 5, 2 or 1. Some of the PAO1 w.t. culture was separated, washed once in M9 buffer, transferred to 1.5 ml plastic micro tubes and heat killed at 65° C for 20 minutes. A sample of the heat killed bacteria was plated on an LB plate to verify inactivation. *Acanthamoeba castellanii* was cultivated, washed and and diluted to 2X10^5^ cells per ml. Twenty seven µL of amoeba suspension were pippeted to all cells of six counting chambers (Vacutest Kima, Italy) and 3 µL of bacterial suspensions were added to final OD_600nm_=0.5,0.2 and 0.1, as well as heat killed w.t. at OD_600nm_=0.5 and no bacteria control (5 replicates per treatment). The number of amoebae cell within the counting grid was counted, this measurement serving as T_0_. The counting cells were kept in a humidified chamber and counted again at 12, 24 and 36 hours.

## Results

Live microscopy of GFP expressing *P. aeruginosa* shows adhesion to *A. castellanii* cells seconds after the introducing bacteria to the amoebae culture (Figure 1).

Quantitative study of *P. aeruginosa* adhesion to the amoebae was conducted using bacterial kinetic adhesion assay in microtiter format[12] (figure 2a), measuring adhesion kinetics in various predator population densities. Initial attachment rates (Figure 2b; first five minutes) are in linear correlation with amoebae population density (R^2^=0.99), while adhesion at one-hour time reaches saturation (Figure 2b). Similar adhesion behavior of *P. aeruginosa* was seen in the presence of paramecium, but not in the presence of nematodes (Figure S1).

To test whether this predator effect on bacterial adhesion kinetics is based on taxis, we followed migration of fluorescent bacteria through a fluorescence blocking 3 μm intervening membrane, in the presence and absence of amoebae in the bottom chamber (Figure 3a), using the Flouroblok™ system (Corning, New York, USA). Figure 3b shows migration was faster in the presence of amoebae. The ability of *P. aeruginosa* to sense amoebae from distance using a soluble moiety is also supported by the modulation of bacterial adhesion behavior by amoebae conditioned medium (Figure 3c).

**Figure 3.**
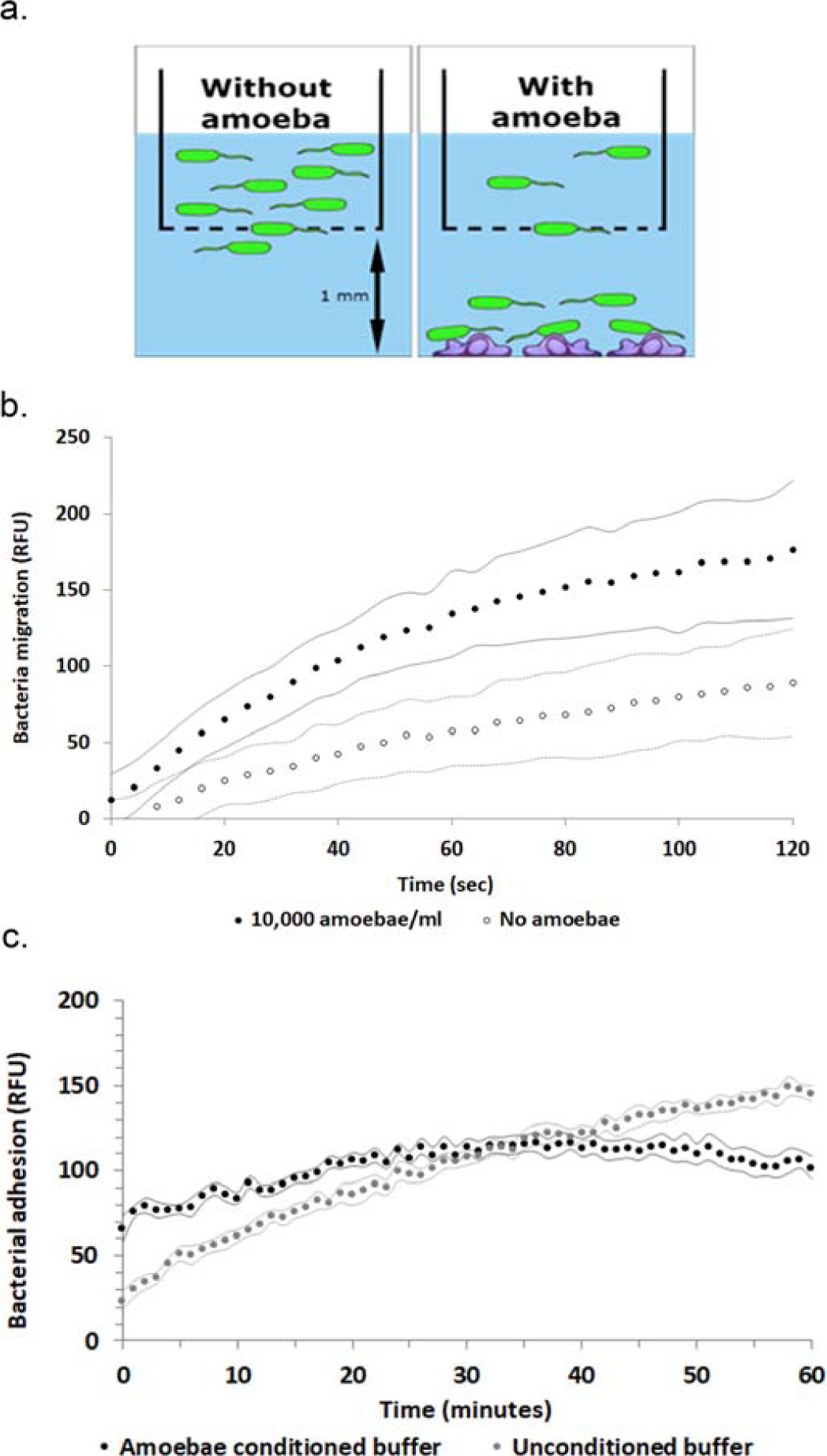
*Pseudomonas aeruginosa* taxis towards amoebae. a. Illustration of measurement of bacterial migration through a fluorescence blocking filter in the absence (left) and presence (right) of amoebae in the bottom chamber. b. Bacterial migration kinetics in the absence (white dots) and presence (black dots) of 10,000 amoebae/ml in the bottom chamber. n=9 for all treatments c. Effect of amoebae conditioned buffer on *P. aeruginosa* adhesion kinetics, n=7. Dots represent measurements, flanking curves stand for ±1 SD.

Mobbing behavior requires an individual not only to sense and a predator, but also to coordinate and synchronize its attack with other individuals, which in bacteria is often facilitated by QS systems. Indeed, *P. aeruginosa* Δ*pqs*A mutant, deficient in PQS production but able to sense and respond to it, exhibits slow adhesion to amoebae cells (Figure 4b). The addition of PQS restores within seconds some of the mutant adhesion behavior, in a dose dependent manner, but only in the presence of amoebae (Figure 4a, 4c). Ten nM of PQS produce a statistically significant increase in attachment within one minute (one tail t test, t_6_=-1.89, p=0.042). Full data set of PQS concentrations is presented in figure S2.

**Figure 4.**
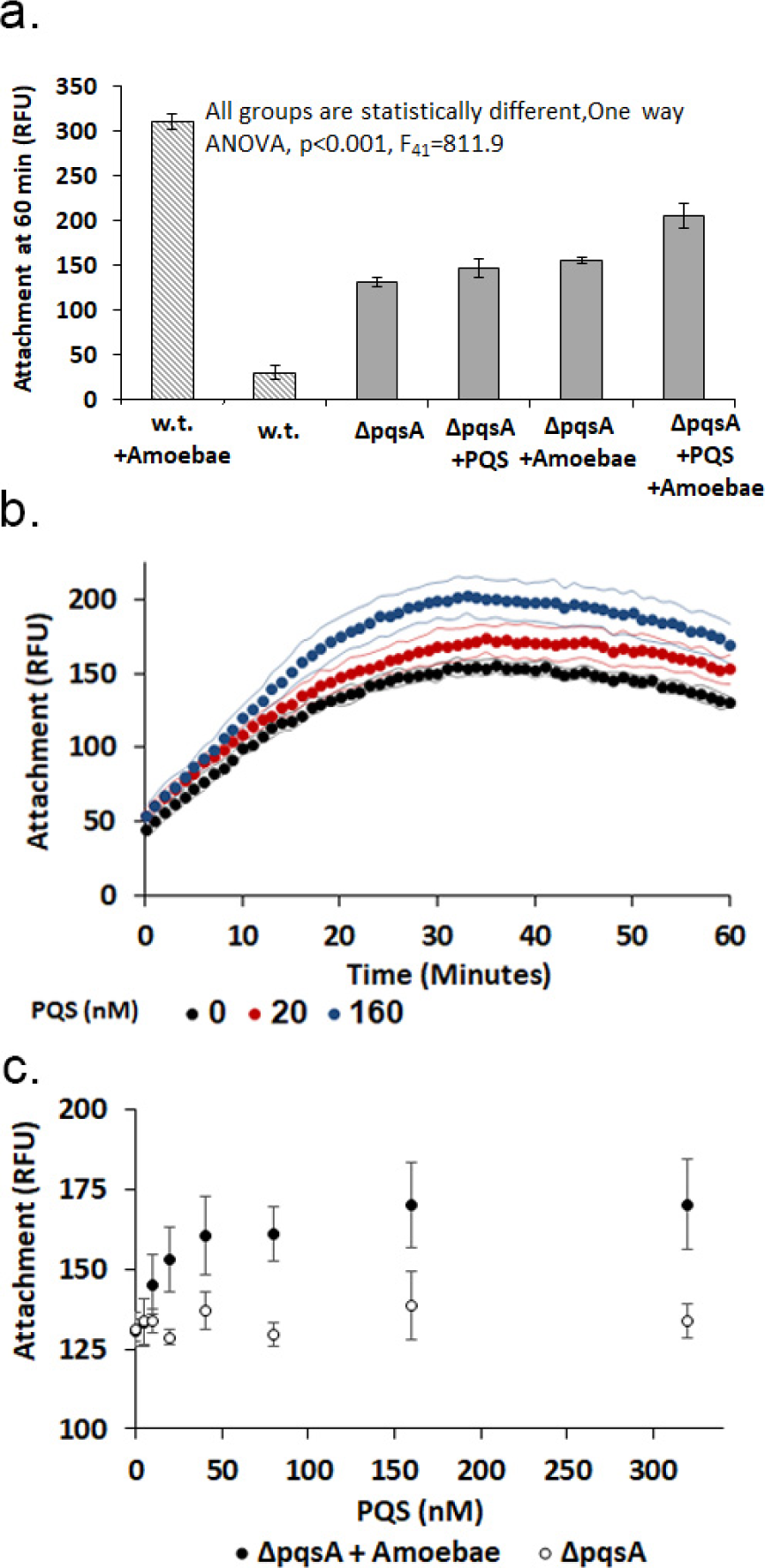
adhesion of *P. aeruginosa* w.t. and ΔpqsA mutant in the presence and absence of amoebae, and with addition of missing PQS: Amoebae treatment consists of 10 000 amoebae per ml (1 000 per well), PQS concentration (when added) is 160 nM. Striped bars stand for w.t., full bars stand for Δ*pqs*A mutant, n=7 for all treatments, error bars represent ±1 SD. **b.** Adhesion kinetics in different PQS concentrations, n=7, dots represent measurements times, flanking curves stand for ±1SD. c. Adhesion at 60 minutes times in the presence (black dots) and absence (white dots) of 10 000 amoebae per ml and in different PQS concentrations. n=7 per treatment, error bars stand for ±1SD.

This immediate effect of PQS signaling on mobbing behavior carries on into hours and days timescales. Amoebae predation affects both w.t. and Δ*pqs*A, but the w.t. population density is reduced by 30% while compared to the mutant which suffered a 55% reduction (Figure 5).

**Figure 5.**
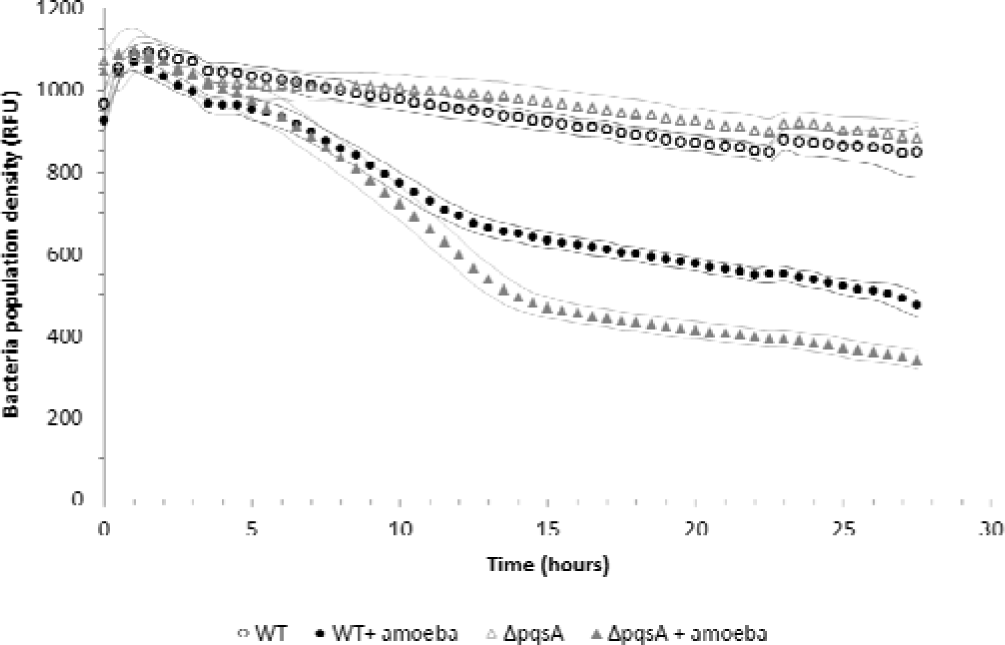
Predation kinetics of *P. aeruginosa* w.t. and Δ*pqs*A. w.t. (black circles) and mutant (grey triangles) fluorescence was measured over 27 hours in the presence (full) and absence (empty) of amoebae. n=14 for all treatments, flanking curves represent ±1SD.

Survival of bacteria corresponded with their ability to kill amoebae (figure 6), studied using direct microscopy counting of amoebae co-cultured with *P. aeruginosa*. Wild type *P. aeruginosa* was able to lyse amoebae while Δ*pqs*A mutant was only able to reduce amoebae growth, when compared to heat killed wild type. Complete dataset, including different initial culture densities, kinetics over three time points, and amoebae growth in co-cultivation with *Escherichia coli* (which enable far better amoebae growth) are found in figure S3.

**Figure 6.**
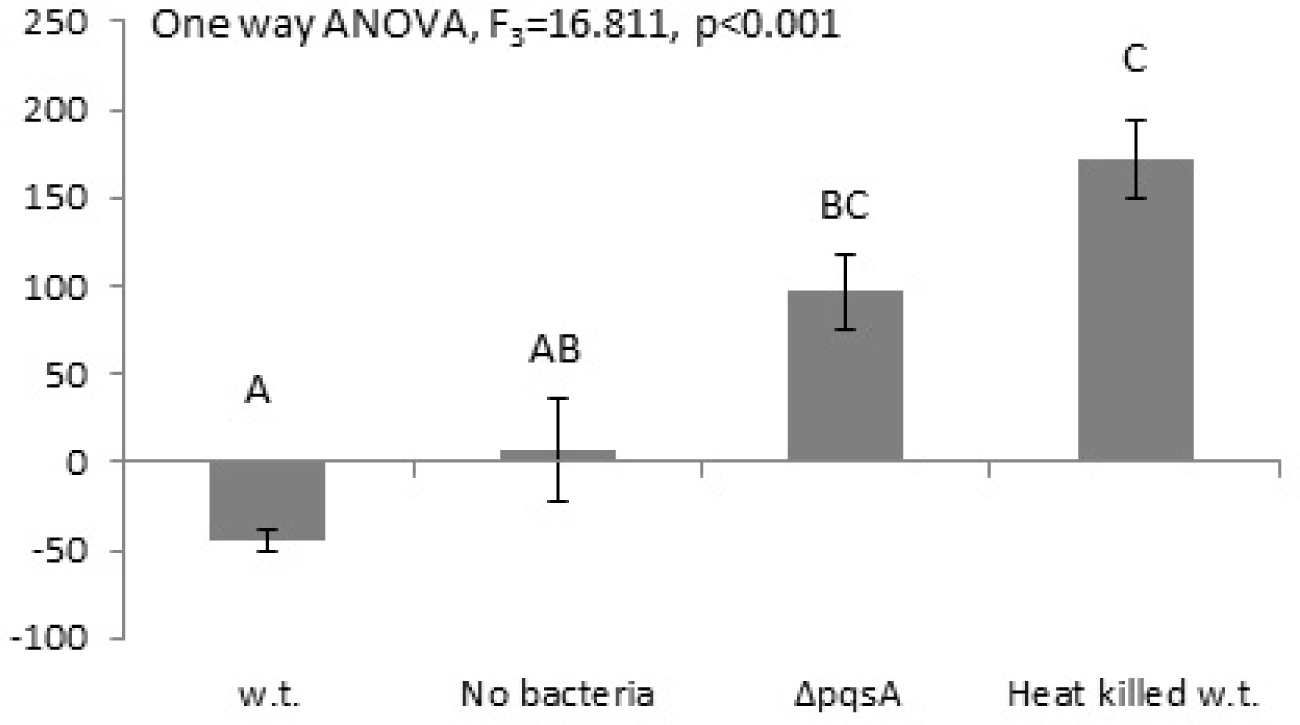
amoebae survival and growth in co-cultivation with w.t. and ΔpqsA *P. aeruginosa*. bars represent % change of initial amoebae count in each chamber at 36 hours’ time. n=5 for each treatment, error bars stand for ±1SD, groups marked with different letters are statistically different.

## Discussion

Mobbing is a predation avoidance behavior, an attack of prey on predator. If enacted by too few individuals such an attack is likely to fail - predators after all evolved by natural selection to deal with prey. Prey can offset this imbalance by a coordinated group attack. The benefit of mobbing; long term reduction in predation risk, is a common good, shared by all members of the prey community. In contrast, the cost of mobbing; immediate predation risk, is paid only by active mobbing participants. This disassociation between benefits and costs reduces the relative fitness of mobbing participants, unless mobbing behavior is prevalent in the prey community. It is not surprising that mobbing behavior is seen in communicating social animal species[13–16], able to generate trust by communicating their willingness to participate in the mobbing effort.

Living is clonal populations that promote kin selection, generating trust by the use of quorum sensing and suffering from predation, mobbing seems a natural course for bacterial evolution. Mobbing, operating in seconds and minutes scale, can buy valuable time, opening the way to slower predation avoidance mechanisms such as formation of micro-colonies or anti-predator toxins.

Time-lapse microscopy of *P. aeruginosa* in the presence of amoebae shows bacterial adhesion to predator cells within seconds of their introduction to amoebae. Bacteria display taxis towards predator cells, which they are able to sense using some soluble predator secretion - a predator kairomone [17].

Coordination of mobbing behavior is seems to be facilitated by the PQS system. A mutant unable to produce PQS was found to be unselective in its adhesion behavior and it ability to kill amoebae. Interestingly, mutants of the LAS and RHL QS systems, both employing N-acyl-homoserine lactone signal molecules, showed w.t. like amoebae killing, suggesting these signals are not involved in mobbing[8]. Given that the PQS is almost unique to *P. aeruginosa*[9] while ASL signals are used by many gram negative bacteria[18], these results demonstrate the importance of a trusted communication, insuring sufficient mobbing participation by competent individuals.

The *P. aeruginosa-A. castellanii* model system described here could be used for the experimental study of behavioral ecology game theory scenarios, enabling easy replication, manipulation and data collection. Microbial ecology is often described only by genetics and metabolism, portraying bacteria as mechanic and passive organisms. This work gives a first impression of bacterial mobbing, a responsive and dynamic behavior. We hope that this, and future of microbes behavioral ecology, may change this view, presenting the true nature of bacteria, as complex and colorful in the micro scale, as our experience of nature in the macro scale. *Quod est inferius est sicut quod est superius* – as above so below.

---

**Supplementary 1.**
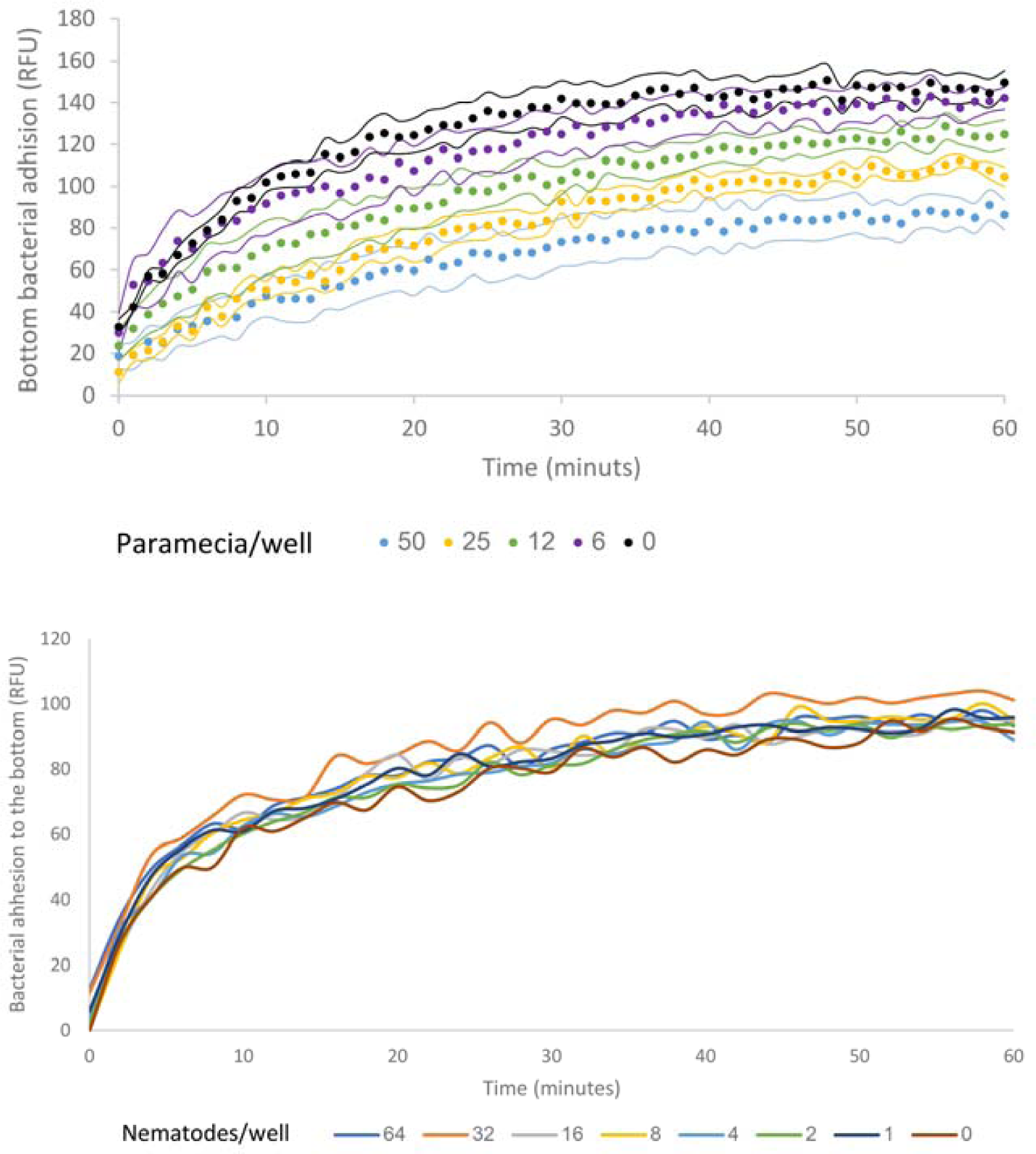
*Pseudomonas aeruginosa* mobbing behavior towards *Paramecia. Sp* and *Caenorhabditis elegans*. Paramecia were separated from wheat grain enrichment culture by filtering, and diluted in de-chlorinated tap water. *C. elegance* were taken from liquid culture and transferred to TBSS. Bacterial suspension used the same medium used in the corresponding predator culture used, added with final concentration of 0.8 mg/ml RED#40. Attachment is measured as bottom fluorescence – as the predators are swimming in the bulk liquid attachment cases reduction in bottom fluorescence kinetics as it removes free bacteria from the medium. Changes in bacterial adhesion are in invers correlation to Paramecium population density but no correlation is seen with nematode population density. n=7 for all treatments, flanking curves (when present) indicate ±1 SD.

**Suplementerary 2.**
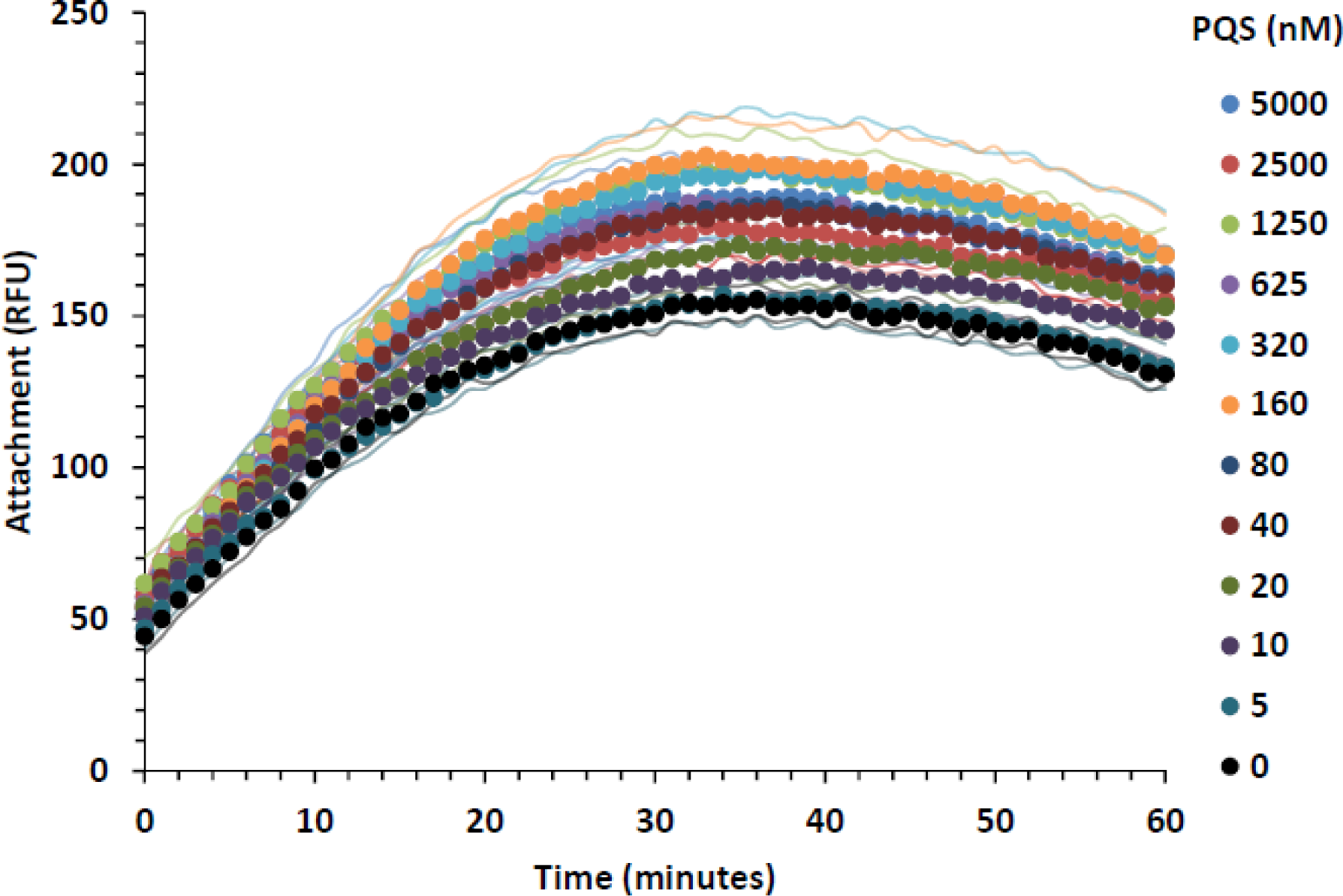
Effect of PQS dose on *P.aeruginosa* ΔpqsA attacment in the presence of amoebae per well: all PQS concentrations presented; n=7; Symbols mark for measurement times; flanking curves signify ± 1 S.D.

**Supplementary 3.**
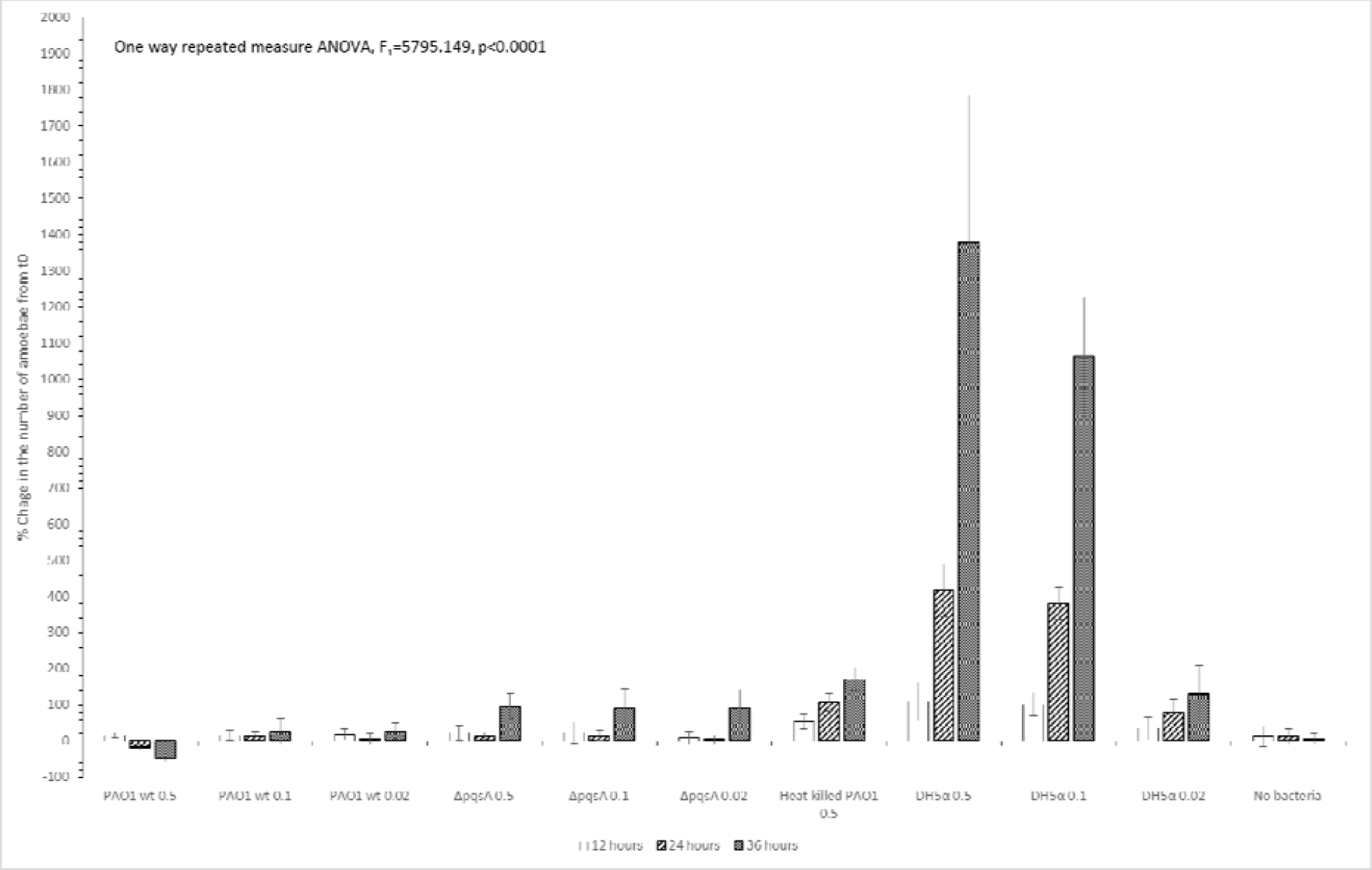
Survival and growth of amoebae in co-cultivation with different bacterial strains, starting with different initial culture densities at three measurement times. n=5 per treatment, bars stand for % change in the number of amoebae cells from time 0, number in category name stand for initial OD_600nm_ error bars represent ±1 SD. All requirements for parametric test were satisfied, Tukey HSD post hoc was applied, results given in the table below.

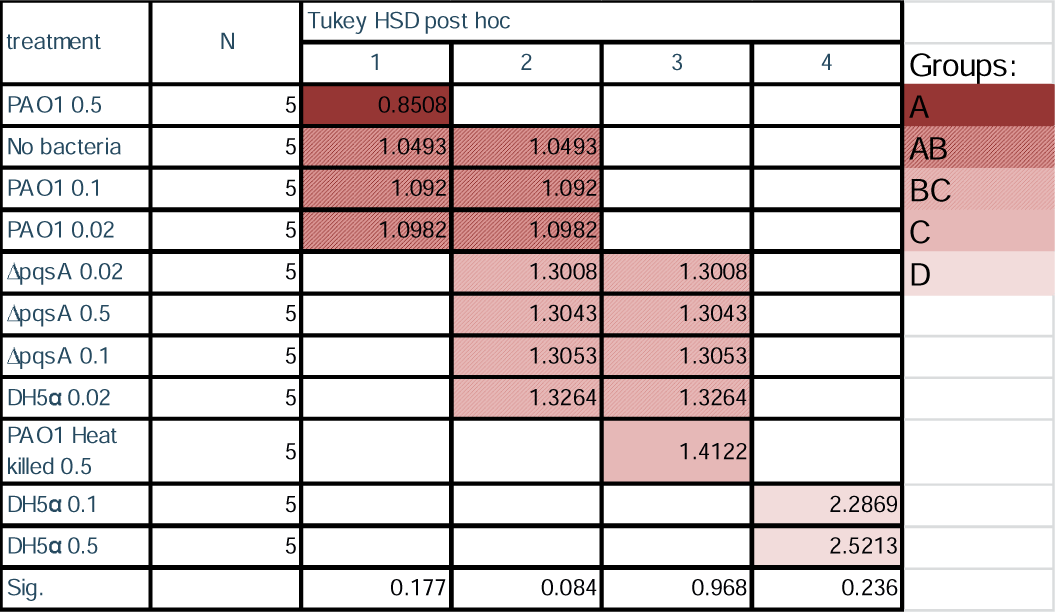

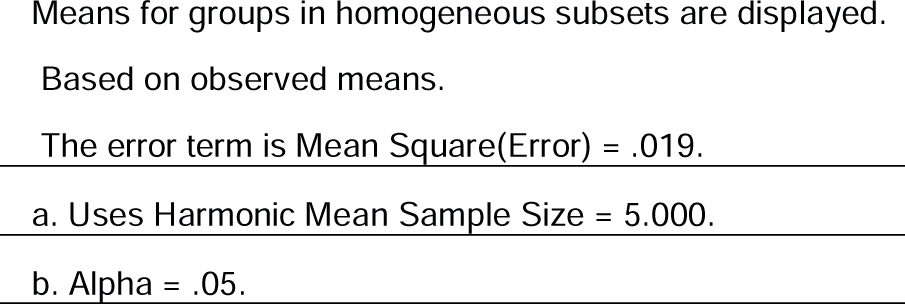

## Notes

### Competing Interest Statement

The authors have declared no competing interest.

